# A novel post hoc method for detecting index switching finds no evidence for increased switching on the Illumina HiSeq X

**DOI:** 10.1101/142356

**Authors:** Gregory L. Owens, Marco Todesco, Emily B. M. Drummond, Sam Yeaman, Loren H. Rieseberg

## Abstract

High throughput sequencing using the Illumina HiSeq platform is a pervasive and critical molecular ecology resource, and has provided the data underlying many recent advances. A recent study has suggested that ‘index switching’, where reads are misattributed to the wrong sample, may be higher in new versions of the HiSeq platform. This has the potential to invalidate both published and in-progress work across the field. Here, we test for evidence of index switching in an exemplar whole genome shotgun dataset sequenced on both the Illumina HiSeq 2500, which should not have the problem, and the Illumina HiSeq X, which may. We leverage unbalanced heterozygotes, which may be produced by index switching, and ask whether the under-sequenced allele is more likely to be found in other samples in the same lane than expected based on the allele frequency. Although we validate the sensitivity of this method using simulations, we find that neither the HiSeq 2500 nor the HiSeq X have evidence of index switching. This suggests that, thankfully, index switching may not be a ubiquitous problem in HiSeq X sequence data. Lastly, we provide scripts for applying our method so that index switching can be tested for in other datasets.

## Introduction

High throughput sequencing, primarily through the Illumina HiSeq platform, has revolutionized molecular ecology. In fact, 50% of original articles in a recent issue of *Molecular Ecology* (Vol 26, Issue 2) included Illumina-derived sequence data. Researchers can now explore questions that were completely unanswerable before current sequencing technologies, using approaches such as genome scans, genome assembly and high density genetic mapping (e.g. Gould and Stinchcombe, 2017; Standage *et al*. 2016; Li *et al*. 2017). With the central role that sequencing plays, it is alarming that a recent preprint suggests increased index switching on the new HiSeq 4000 and HiSeq X machines (Sinha *et al*. 2017).

To prepare DNA for Illumina sequencing, strands are fragmented and adapter sequences are attached to the ends of these fragments. These adapters contain the sequence that binds to the flow cell, a primer sequence for amplification during sequencing and, potentially, a barcode index for linking reads to individual samples. Indexes are required when multiplexing samples within a single sequencing lane, and can be included in adapters at one or both ends of the DNA fragments. As the output of a single sequencing lane has increased, multiplexing has become increasingly common. This is especially true in molecular ecology, where researchers often aim to maximize sample size by using low coverage whole genome data (Buerkle and Gompert 2013). For example, a single lane on the HiSeq 4000 can sequence 200 stickleback genomes (~460MB) to 1× coverage. Consequently, it is critical that samples are correctly demultiplexed or the resulting sequence data will contain mixes of reads from unexpected and unpredictable sources.

A recent preprint by Sinha *et al*. reports high levels of index switching in a single cell RNAseq experiment (Sinha *et al*. 2017). They dual indexed (i.e. barcodes on both adapters) all samples using a Nextera XT kit and found that samples that shared a single index had greater similarity in gene expression levels than expected. The authors attributed this to index switching, and showed that controls containing adapters and index primers but no template DNA still had reads assigned to them, receiving 5–7% of the average number of reads of samples with template DNA as a result of index switching. They proposed that index switching occurs during cluster generation (before sequencing) when free index primers replicate already indexed library fragments. These newly copied fragments will then carry one wrong index and be misattributed to another sample. Importantly, they find that this only occurs on the Illumina HiSeq 4000, which uses a patterned flow cell and a new exclusion amplification (ExAmp) chemistry, and not in the NextSeq 500, which does not. Both the HiSeq 4000 and HiSeq X use a patterned flow cell and the cBot 2 system for cluster generation, suggesting that the problem may occur in both machines. Illumina has acknowledged that index switching can occur and is higher in machines that use a patterned flow cell, but suggests total index switching is >2% of reads (Illumina, 2017).

In light of the potential problems, we explored a set of whole genome sequenced samples, half of which were sequenced on the HiSeq 2500, which does not use the patterned flow cell and ExAmp chemistry, and half on the HiSeq X, which does. We have developed a novel method for detecting index switching in genomic datasets and show that in our samples index switching is minimal and not enriched in the HiSeq X.

## Methods

### Study species and library preparation

To identify whether index switching was detectable in an average whole genome sequence dataset, we analyzed a set of 323 wild *Helianthus annuus* (common sunflower) whole genome sequence samples. Plants were grown from field-collected seeds obtained from 28 populations located across the Midwestern USA and Southern Canada. Genomic DNA was extracted from frozen leaf tissue using either a modified CTAB protocol (based on Murray and Thompson, 1980), the DNeasy Plant Mini Kit or a DNeasy 96 Plant Kit (Qiagen, Hilden, Germany). DNA was sheared to an average fragment size of 350 bp using a Covaris M220 ultrasonicator (Covaris, Woburn, Massachusetts, USA), following the manufacturer’s recommendations. 750 ng of sheared DNA were used as starting material to prepare paired-end whole-genome shotgun Illumina libraries, using a protocol largely based on Rowan *et al*, 2015, the TruSeq DNA Sample Preparation Guide from Illumina (Illumina, San Diego, CA, USA) and Rohland and Reich, 2012. End-repairing of the sheared DNA fragments was performed using the NEBNext End Repair Module (NEB, Ipswich, Massachusetts, USA). The fragments were then A-tailed using Klenow Fragment (3’-->5’exo-) from NEB and ligated to 24-bp-long, non-barcoded adapters with a 3’ T-overhang (Table S1) using the Quick Ligation Kit from NEB. After each enzymatic step, the reactions were purified using 1.6 volumes of a solution of paramagnetic SPRI beads (MagNA), prepared according to Rohland and Reich, 2012. An enrichment step was then performed using KAPA HiFi HotStart ReadyMix (Roche, Basel, Switzerland) and short, non-indexed primers that do not extend the adapters (Table S1). The reactions were then purified using 1.6 volumes of MagNA beads. The sunflower genome contains a very large amount of highly repetitive sequences derived from the recent expansion of two retrotransposon families (Staton *et al*. 2012). In order to reduce the representation of repetitive sequences, the enriched libraries were treated with a Duplex-Specific Nuclease (DSN; Evrogen, Moscow, Russia), following the protocols reported in Shagina *et al*. 2010 and Matvienko *et al*. 2013, with modifications. The fragments were then further amplified using Kapa HiFi HotStart ReadyMix and primers (to a final concentration of 0.4 μM each) to complete the adapters and add a six-bp index to the P7 adapter (Table S1). The sequence of the completed adapters is identical to that Illumina’s TruSeq adapters.

After amplification, the libraries were purified twice with 1.6 volumes of MagNA beads, quantified using a QuBit dsDNA Broad Range Assay Kit (Invitrogen, Carlsbad, California, USA) and analyzed on a 2100 Bioanalyzer instrument using a High Sensitivity DNA Analysis Kit (Agilent, Santa Clara, California, USA). The libraries were then quantified on an iQ5 Real Time PCR Detection System (Bio-Rad, Hercules, California, USA) using Maxima SYBR Green qPCR Master Mix (ThermoFisher Scientific, Waltham, Massachusetts, USA) to determine molarity, and pools consisting of ten libraries each were prepared. All libraries were sequenced at the Genome Quebéc Innovation Center; 156 libraries were sequenced on a HiSeq 2500 instrument and 165 were sequenced on a HiSeq X instrument (Illumina, San Diego, CA, USA). Importantly, samples were multiplexed within lanes in a random manner without regard to population ID.

### Variant calling

We aligned all samples to the *H. annuus* XRQ genome using BWA (version 0.7.9a), removed PCR duplicates using samtools and called variants using FreeBayes (version 1.1.0) (Li and Durbin 2010; Li *et al.*, 2009; Garrison and Marth 2012). In all cases, we used default parameters. For this analysis, we selected di-allelic SNPs with QUAL > 30 using vcflib (https://github.com/ekg/vcflib).

### Testing for index switching

To identify whether index switching is increased in samples sequenced on the HiSeq X, we leveraged the fact that individual samples in our dataset were either sequenced on the HiSeq X or the HiSeq 2500. Therefore, we can not only estimate index switching rates on the HiSeq X, but also tell if it is higher than for previous technology.

Previous work has suggested that index switching is occurring for 1–10% of reads depending on factors during library preparation and sequencing (Sinha *et al*. 2017). This low level means that, for our dataset, at a single locus, an allele acquired because of index switching is likely to only have one read, given moderate overall read depth. We looked for these unbalanced heterozygotes (i.e. one read for allele 1, many reads for allele 2) and asked if the rare allele (i.e.. the under-sequenced allele) was found in other samples sequenced in the same lane (which we refer to as “allele sharing”). We then calculated 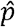, the probability that the rare allele should be found in those samples based on *f*, the allele frequency for all samples sequenced with that machine (excluding the unbalanced focal individual) and *n*, the number of other samples with genotypes in the lane (1).

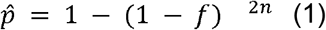

We then plotted 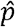, the predicted proportion of cases where the allele is present in at least one copy in the other samples from the lane, against *p*, the observed proportion of cases with allele sharing. We fit a line to this relationship using a generalized additive model in the *stat_smooth* command from *ggplot2* (Wickham, 2016). If index switching is not occurring, we expect a straight line at 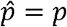. Alternatively, if index switching is occurring, we expect 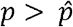 indicating greater sharing of under-sequenced alleles within a lane than expected by chance. These proportions were calculated independently for HiSeq 2500 and HiSeq X samples, using the first 500,000 variable sites in the genome.

As a control, for each unbalanced heterozygote we calculated the *p* using the same number of genotyped samples sequenced using the same machine, but not the same lane. This control should not show evidence of index switching.

It's important to note that if samples were sorted into sequencing lanes based on a genetic grouping (e.g. species or population), we would find 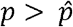 in the absence of index switching. In our dataset this is not the case, as samples were randomly assigned into lanes.

### Simulations

To explore the sensitivity of our measure of index switching, we bioinformatically switched reads in our vcf file, randomly selecting 0, 0.1, 0.5, 1, 5, or 10 percent of reads at each site across all individuals to be switched. Switched reads were removed from the individual (i.e. reducing read depth) and added to another individual sequenced in the same lane (i.e. increasing read depth). We then recalculated genotypes simply by assigning samples containing reads for both alleles as heterozygotes. These simulations were run through the same algorithm to detect index switching.

### Data availability

All scripts used in this manuscript are available on github (<u>https://github.com/owensgl/index_investigator</u>) along with a dataset containing lane identifiers, genotypes and read depths for samples used in this study.

## Results

We fail to find evidence that index switching is occurring in our dataset. For samples sequenced on both machines, the observed proportion of allele sharing within a lane tracked the predicted proportion closely (Figure 1, Supplementary figure 1). This was consistent with the pattern seen in our control that used samples from different lanes. Despite this, we find that our method is able to identify index switching in the simulated dataset. In particular, we find elevated allele sharing around 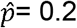, even when index switching only represents 1% of reads (Figure 2). In our dataset, 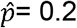 corresponds to rare alleles (minor allele frequency < 5%). This makes sense because common alleles are expected to have high allele sharing even in the absence of index switching which makes the signal more difficult to observe.

**Figure 1.**
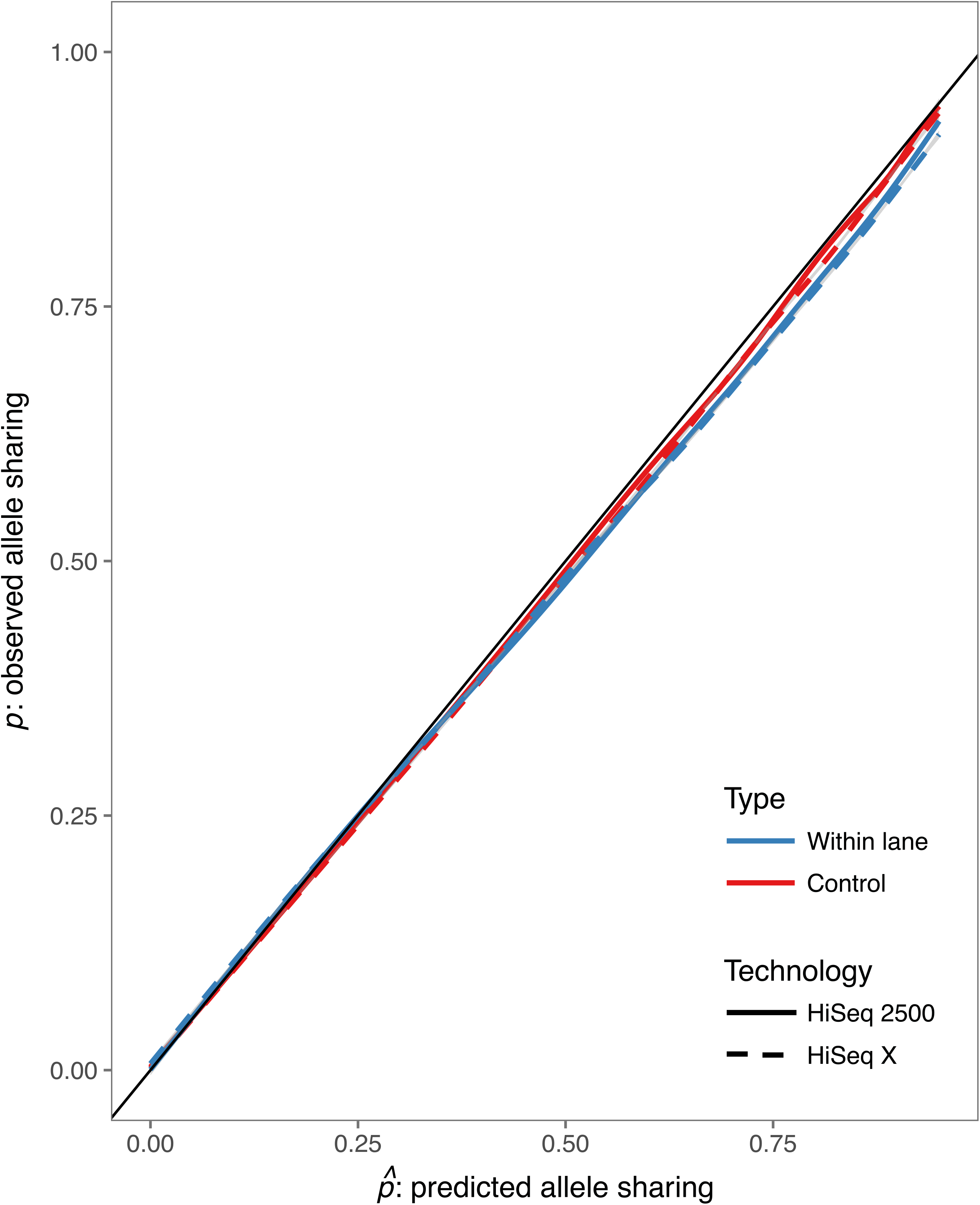
(a) The relationship between predicted allele sharing and observed allele sharing for samples sequenced on the HiSeq 2500 (solid line) and HiSeq X (dashed line). Allele sharing was calculated for samples sequenced together in a lane (blue) and for a control group sequenced in different lanes (red).

**Figure 2.**
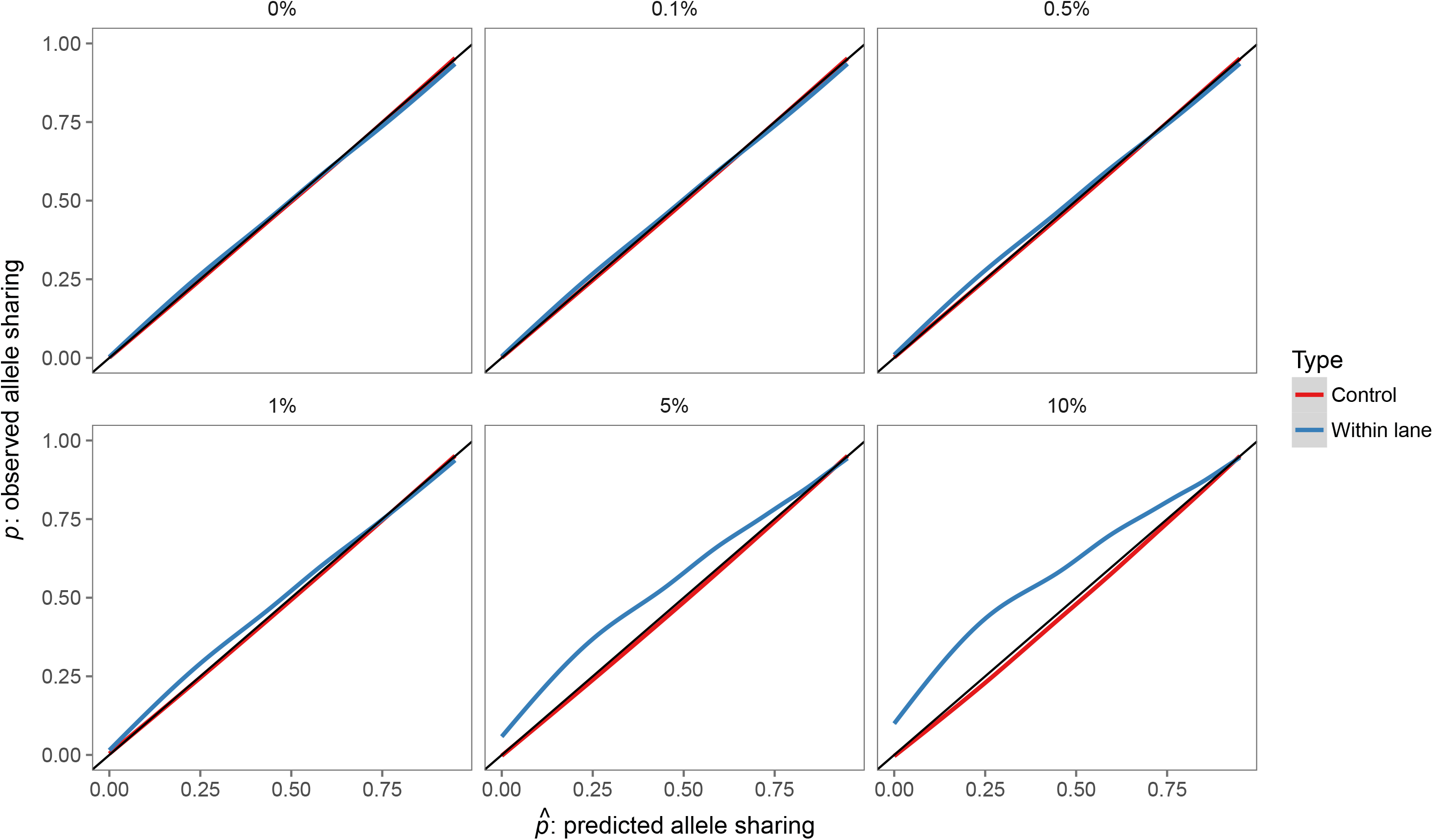
The relationship between predicted allele sharing and observed allele sharing with different degrees of simulated index switching. The 0% index switching test controls for the re-calling of genotypes that occurs during simulated index switching. Allele sharing was calculated for samples sequenced together in a lane (blue) and for a control group sequenced in different lanes (red).

## Discussion

Widespread, undetected index switching represents a nightmare scenario for molecular ecologists worldwide. Here we show that in one exemplar dataset, index switching is not higher in samples sequenced on the new patterned flowcells and is likely below 1% of reads. Furthermore, we provide a way to visualize index switching for sequenced genomic datasets.

### Why don’t we find index switching?

Our results are clearly different from Sinha *et al*., who found index switching affecting 5–10% of reads. This could potentially be caused by differences in sequencing library preparation. Sinha *et al*. used cDNA as starting material and the Nextera tagmentation technology from Illumina to fragment the DNA and tag the fragments with adapters, whereas we used genomic DNA sheared using ultrasonication and then added the adapters to the fragments via enzymatic ligation. Furthermore, our protocol included a depletion step, to reduce repetitive elements in the genome, that is not present in the Nextera XT protocol. However, the final step of library preparation is substantially equivalent between the two approaches; DNA fragments with short adapters at their extremities are PCR-amplified using primers that complete the adapters and add unique sequence indices, allowing pooling of different samples in a single flow cell. Given that carry-over of free indexed primers from this step is the likely cause of index switching during the ExAmp procedure (Sinha *et al*. 2017), the two approaches can be confidently compared for the purpose of investigating the occurrence of index switching.

Another possible difference between the two experiments is that, while the Nextera XT kit uses dual indices (i.e. both the P5 and P7 adapters are indexed), we used only a single index on the P7 adapter. This has the potential to halve index switching in our dataset, assuming that switching occurs equally from both adapters. If the unindexed P5 adapter were to be replaced in our dataset, this would not result in index switching because no index is present. For a dual indexed library, it would result in index switching.

Finally, the main difference we noticed between our libraries and the one shown in Figure 4B of Sinha *et al*. is the large amount of free adapters/primers that are found in the latter (compare with the Bioanalyzer plot for one of our libraries in Figure 3a). Our enhanced cleanup efficiency could be due to fact that, while the Nextera XT kit recommends a single cleanup step with 0.6 volumes of Agencourt AMPure XP beads, we performed two rounds of cleanup with 1.6 volumes of MagNA beads (the maximum size of the fragments that are removed during beads cleanup is, roughly, inversely proportional to the ratio of bead solution that is added to the reaction - smaller volumes of beads should therefore be more efficient at removing free adapter/primers). However, a single cleanup with 1 volume of MagNA beads was sufficient to completely remove primers/adapters from our libraries (Figure 3b). MagNA and AMPure XP beads have been shown to have comparable recovery efficiency and size discrimination (Rohland and Reich, 2012), and this is confirmed by our experience. While it is possible that, because of their different design, libraries produced using the Nextera XT protocol simply contain a much larger amount of free adapters/primers that cannot be efficiently removed with one single cleanup step, we did not directly test this.

**Figure 3.**
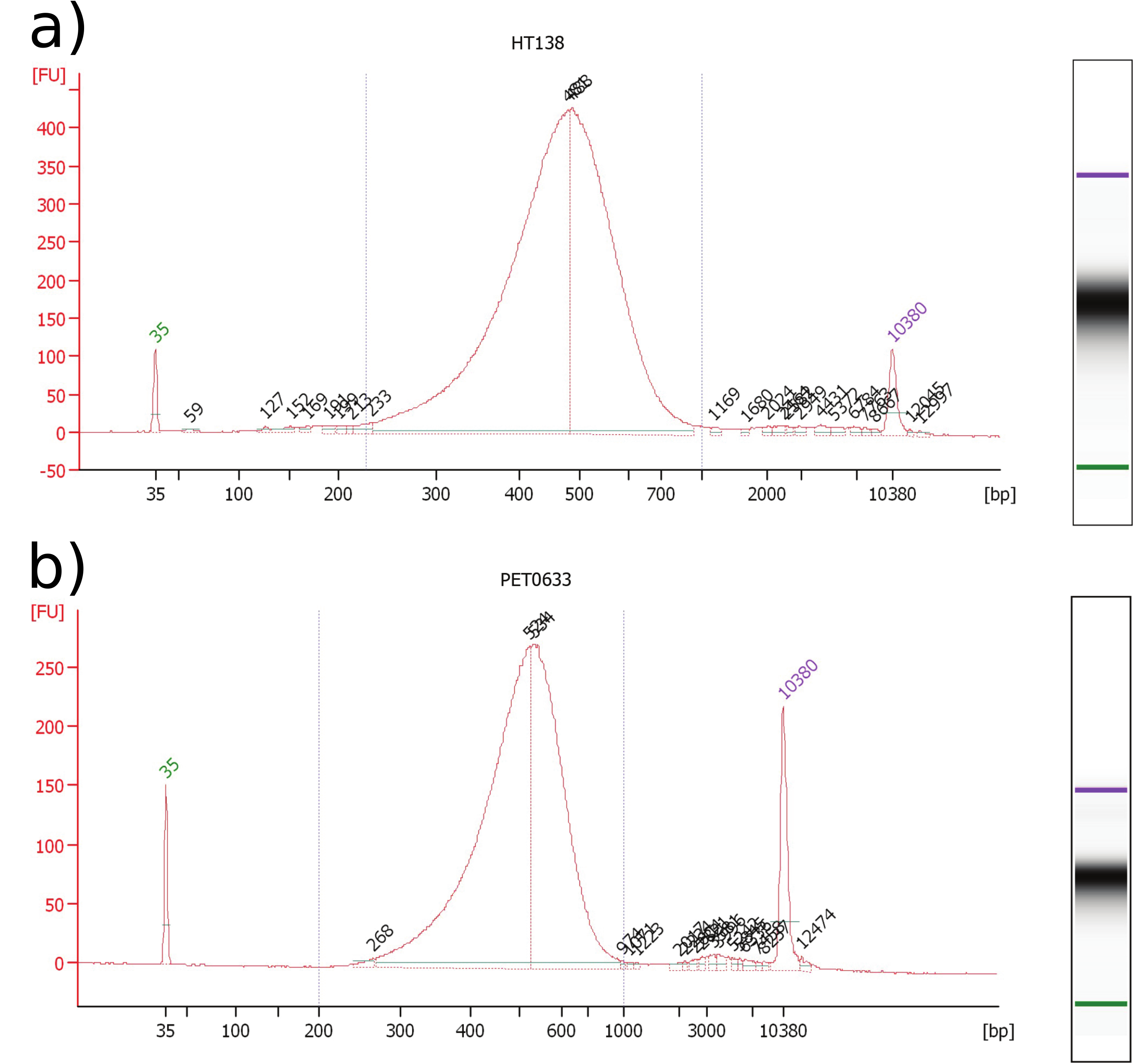
Bioanalyzer plots for representative whole genome shotgun sequencing libraries used in this study, after the final amplification and cleanup step. The plot shows the abundance of fragments of different sizes in the library (measured in fluorescence units, FU). The peaks at 35 bp (green) and 10,380 bp (purple) are internal standards. Free index primers should appear as a peak at ~50 bp. a) Library that underwent two rounds of cleanup after PCR amplification, each using 1.6 volumes of MagNA beads. b) Library that underwent a single round of cleanup with 1 volume of MagNA beads.

### When is index switching confounding?

Certain kinds of experiments are more likely to be affected index switching. Gene expression quantification using RNAseq is especially sensitive because highly expressed genes can bleed into other samples, homogenizing expression measures with lanes. In cancer genomics, low frequency alleles represented by a minority of reads are both important and can be produced by index switching. Similar issues can occur in Pool-seq experiments used in molecular ecology, where index switching could affect estimation of allele frequencies, slightly homogenizing differences among pools sequenced in the same lane.

For high coverage genomic sequencing of diploid organisms, index switching can produce unbalanced heterozygotes, where one allele is represented by one or two reads and the other by many reads. These present a genotyping challenge because unbalanced heterozygotes can also be produced naturally by stochastic sampling of alleles or via PCR bias during library prep. Future genotyping programs may use haplotype information of reads along with sequencing lane identity to detect when index switching is occurring and remove contaminants. In low-coverage genome sequencing, identifying individual instances of index switching may be impossible and will result in an increased rate of false heterozygote genotype calls (when an index is switched among alternate homozygotes) and slightly increased quality scores for heterozygotes mis-called as homozygotes (when all sequenced reads represent only one allele of the heterozygote and an index is switched from a homozygote with the same allele).

Which samples are multiplexed in a lane has a large effect on whether index switching is a problem. If each sample represents a distinct, distantly related species, then misattributed reads are unlikely to align to a reference genome. If all samples are from a single population, misattributed reads are more likely to carry alleles already present. In the worst case scenario, samples of closely related species or distantly related populations with misattributed reads could be mistakenly inferred as novel alleles. This could reduce divergence estimates like F_ST_ or confuse phylogenetic signals. Although stringent allelic balance cut offs for heterozygous genotypes would remove the false heterozygotes from index switching, it may also remove true heterozygotes or miscall them as homozygotes, especially at lower (<10) average read depth.

### Best practices to avoid index switching

Although we failed to detect index switching here, it may be prudent to employ techniques for avoiding the issue. Two main suggestions have been proposed: (1) using dual index barcodes, so that both indices are unique to a sample and (2) thoroughly cleaning library preparations to remove free primers. Beyond this, researchers should be more aware of what samples are multiplexed together, a process that is often determined by the sequencing facility without regard for sample identity.

## Conclusion

We have failed to find evidence for index switching here, but we certainly do not make the claim that it cannot or does not happen. However, we would like to make two points: (1) index switching does not always occur and (2) that vigilance is necessary. With greater attention to this problem, research labs and companies can spend time and effort creating molecular protocols to reduce this issue and bioinformatic programs to detect or remove misattributed reads. Like all genotyping methods, errors are inevitable, but by better understanding their source we can sort signal from noise.

## Figures

Supplementary Figure 1. A stacked histogram of the allele sharing presence/absence at different 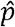 values for within lane samples (a) and control samples (b). This is the raw data used to produce Figure 1. The difference in heights between HiSeq 2500 and HiSeq X reflects differences in sequencing depth and multiplex pooling. The HiSeq X produced more reads, less missing data and more unbalanced heterozygotes.

